# A sustained Hox program delineates brainstem neurons essential for breathing

**DOI:** 10.1101/2025.08.01.668179

**Authors:** Matthew T. Moore, Minshan Lin, Alicia N. Vagnozzi, Raquel López de Boer, Elyse M. Brozost, Lucie Jeannotte, Lindsay A. Schwarz, Polyxeni Philippidou

## Abstract

Respiratory neurons in the brainstem must diversify and acquire unique properties during development to enable breathing at birth. Dbx1-expressing progenitors give rise to functionally and molecularly distinct excitatory respiratory populations, including rhythm-generating pre-Bötzinger complex (preBötC) neurons and phrenic motor neuron (MN)-projecting rostral Ventral Respiratory Group (rVRG) neurons. These neurons are organized rostrocaudally within the ventral respiratory column (VRC) but, despite their critical functions in breathing, the mechanisms that control their organization and diversification are not well understood. Here, we generate a novel genetic tool to label brainstem neurons within the VRC. We find that rVRG neurons selectively express *Hox5* genes through postnatal stages. Selective deletion of all *Hox5* paralogs from *Dbx1*-derived neurons leads to respiratory dysfunction, perinatal death, and changes in the pattern and amplitude of phrenic MN firing. We show that *Hox5* inactivation leads to a caudal expansion of putative preBötC neurons, likely at the expense of the rVRG. Collectively, our findings indicate that Hox5 proteins are required for the delineation and functional specialization of excitatory brainstem neurons essential for breathing.

## Introduction

The neural circuits that control breathing must assemble with high fidelity during development to generate efficient breathing upon birth. The rostrocaudally-arrayed ventral respiratory column (VRC) in the brainstem contains distinct neuronal populations essential for breathing^1,2^. Pre-Bötzinger complex (preBötC) neurons generate the rhythmic pattern of breathing and project to rostral Ventral Respiratory Group (rVRG) neurons that activate phrenic motor neurons (MNs) in the spinal cord to drive diaphragm muscle contractions^3^. To support their unique functions, VRC neurons must acquire different properties. For example, preBötC neurons must form recurrent connections largely within the brainstem that enable them to generate the breathing rhythm^4-7^, while rVRG neurons are non-rhythmogenic and send long distance projections that target phrenic MNs in the spinal cord^8,9^. Within the preBötC, further diversification of neuronal subtypes, indicated by the expression of specific markers such as somatostatin (SST)^10-13^, reelin^14^, and neurokinin 1 receptor (NK1R)^15,16^, enables highly specialized functions in rhythm and pattern generation^6,17-19^. While we lack this level of granularity for pre-motor regions such as the rVRG, largely due to the absence of known specific molecular markers, it is likely that considerable heterogeneity exists within this population as well. Despite stark distinctions in their anatomy and function, excitatory neurons within both the preBötC and rVRG are generated from the same progenitor domain, which expresses the homeobox transcription factor Dbx1^10,11,20,21^. This indicates that additional developmental programs, independent of their progenitor identity, must tightly regulate their firing patterns and connectivity, as well as ensure that the correct subtypes are generated in the appropriate location and proportion. To date, we have a limited understanding of how distinct respiratory neuron subtypes acquire unique properties that enable their specific functions throughout the respiratory circuit.

Mutations in several genes encoding transcription factors lead to disruptions or cessation of the breathing rhythm and perinatal lethality. These can be largely classified into mutations that disrupt rhombomere segmentation and alter brainstem organization during early hindbrain development^22-29^, those resulting in the agenesis of specific respiratory populations^30-32^, or those that broadly affect several classes of respiratory neurons. For example, deletion of *Dbx1*, expressed in the domain that gives rise to nearly all glutamatergic neurons in the VRC, halts the respiratory rhythm^10,11,21^. Deletions of certain *Hox* genes, such as *Hoxa1* and *Hoxa2*, also lead to early developmental defects and disrupted breathing, but these effects are likely due to their impact on more rostral breathing structures, such as pontine nuclei and the parafacial respiratory group (pFRG), rather than more caudal VRC populations^33-38^. While Hox proteins, such as Hox4 paralogs, are highly enriched in postmitotic VRC neurons up to postnatal stages, their cell-autonomous functions in these neurons are not known^39-41^.

Here, we develop a novel genetic mouse tool to delineate VRC neurons during development and early postnatal stages. We find that Hox5 proteins are selectively and continuously expressed in rVRG but not preBötC neurons. Selective *Hox5* deletion from *Dbx1*-derived neurons leads to perinatal lethality and a dramatic reduction and disorganization of phrenic MN activity. Analysis of brainstem respiratory populations shows a significant caudal expansion of SST+ putative preBötC neurons, likely at the expense of rVRG pre-motor neurons. Altogether, our data indicate that Hox5 proteins are required for the delineation and functional specialization of the brainstem neurons that underlie breathing.

## Results

### A novel molecular tool to target the ventral respiratory column

rVRG neurons provide the major excitatory input to phrenic MNs and are a critical node in the inspiratory circuit, as they integrate inputs from several key respiratory areas in the brainstem to coordinate the activity of cranial and phrenic MNs^8,42,43^. While rVRG neurons can be defined anatomically using landmarks such as local motor nuclei^44-46^, the molecular programs that control their development, connectivity, and function are mostly unknown, largely due to a lack of specific tools to label and target the rVRG. We previously identified the cell adhesion molecule Cadherin 9 (Cdh9) as a putative marker of phrenic MN-targeting rVRG neurons (Fig. 1A)^47^. To test whether Cdh9 could serve as a novel rVRG molecular marker, we generated a *Cdh9::FlpO* mouse using CRISPR-Cas9 technology. Briefly, a *P2A-FlpO* sequence was inserted in place of the stop codon in *Cdh9* exon 12 (see Materials and Methods). We crossed *Cdh9::FlpO* mice to *RCE:FRT* reporter mice^48^ to express Flp-dependent GFP in *Cdh9*-expressing populations (referred to as *Cdh9*^*GFP*^).

**Figure 1.**
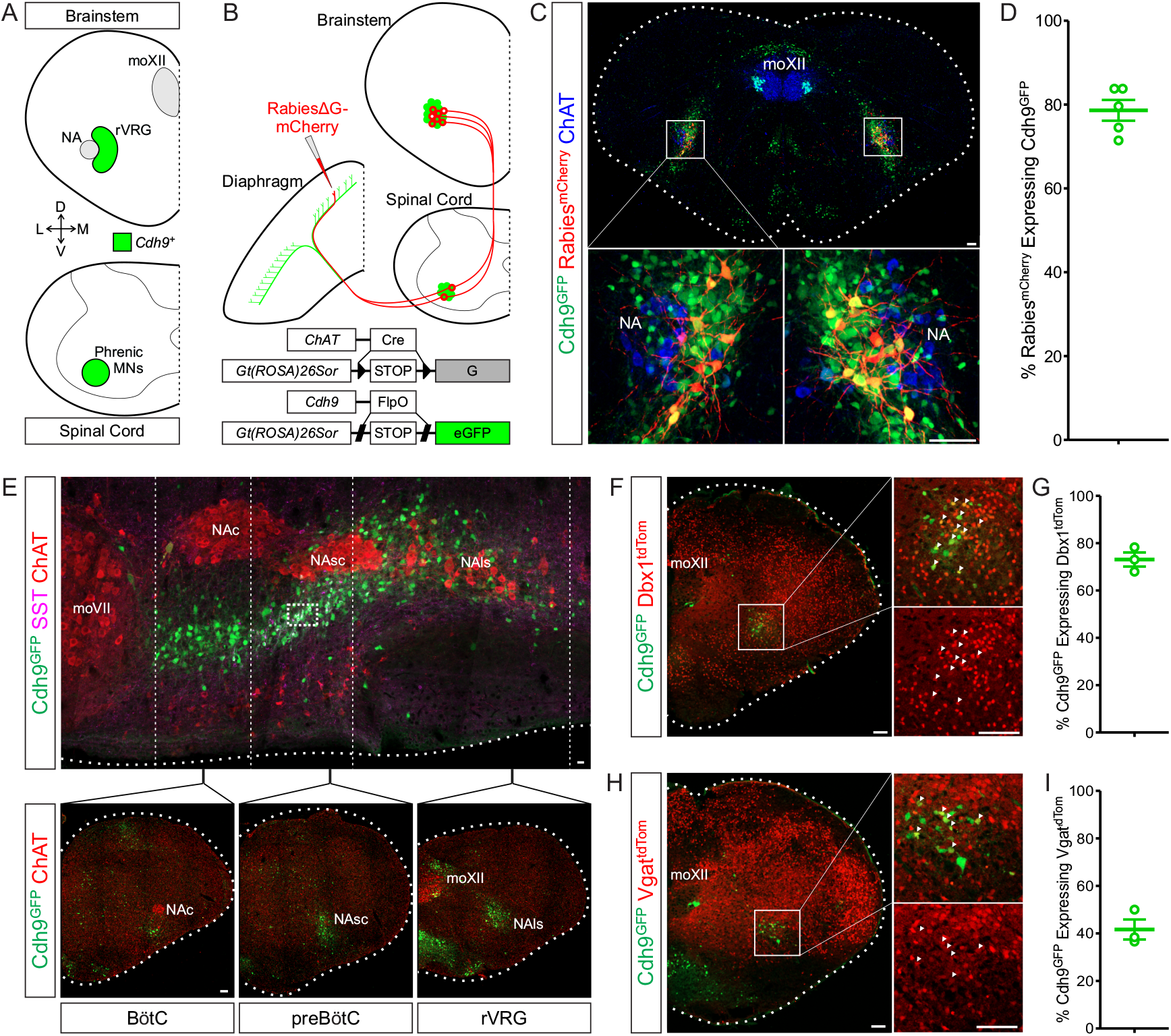
A novel molecular tool targets the ventral respiratory column. (A) Diagram of *Cadherin 9* (*Cdh9*, green) expression in the developing respiratory neural circuit. In the brainstem (top), *Cdh9* is expressed in the rostral Ventral Respiratory Group (rVRG) adjacent to the nucleus ambiguus (NA) at the level of the hypoglossal motor nucleus (moXII)^47^. In the spinal cord (bottom), phrenic motor neurons (MNs) also express *Cdh9*^63^. (B) Schematic of rabies-mediated retrograde transsynaptic tracing strategy. A glycoprotein (G)-deficient rabies virus (RabiesΔG-mCherry) is injected into the diaphragm of *ChAT::Cre; RphiGT; Cdh9::FlpO; FRT-GFP* mice at P4. (C) Example of phrenic MN-targeting rVRG neurons, labeled by mCherry (red), expressing *Cdh9::Flp*-dependent GFP (Cdh9^GFP^, green) at P11. MN populations are labeled by choline acetyltransferase (ChAT, blue). Scale bar = 100 μm. (D) Percentage of mCherry+ cells expressing Cdh9^GFP^ at P11 (78.6 ± 2.5% of mCherry+ neurons, n = 5). (E) Sagittal section through the VRC of a *Cdh9*^*GFP*^ mouse at P14. ChAT (red) labels the facial motor nucleus (moVII) and the compact (NAc), semi-compact (NAsc), and loose (NAls) partitions of the NA. Somatostatin (SST, magenta-inside dashed line box) delineates the pre-Bötzinger Complex (preBötC). Coronal sections (below) show the mediolateral distribution of Cdh9^GFP^ (green) neurons. Scale bar = 100 μm. (F) *Dbx1::Cre*-mediated tdTomato (Dbx1^tdTom^, red) and Cdh9^GFP^ (green) expression in the rVRG at P0. Scale bar = 100 μm. (G) Percentage of Cdh9^GFP^ neurons expressing Dbx1^tdTom^ at P0 (73.1 ± 3.0% of Cdh9^GFP^ neurons, n = 3) (H) *Vgat::Cre*-mediated tdTomato (Vgat^tdTom^, red) and Cdh9^GFP^ (green) expression in the rVRG at P0. Scale bar = 100 μm. (I) Percentage of Cdh9^GFP^ neurons expressing Vgat^tdTom^ at P0 (41.6 ± 4.2% of Cdh9^GFP^ neurons, n = 3)

To determine whether *Cdh9*^*GFP*^ mice can be used as a molecular tool to label phrenic MN-targeting rVRG neurons, we performed rabies virus-mediated monosynaptic retrograde tracing to visualize inputs to phrenic MNs. We utilized a modified glycoprotein-depleted mCherry-tagged rabies (RabiesΔG-mCherry) virus, which requires expression of glycoprotein G for transsynaptic transport and therefore cannot be transsynaptically transported in mammalian neurons. We crossed *RphiGT* mice, which express glycoprotein G after Cre-mediated recombination, to *choline acetyltransferase* (*ChAT)::Cre* mice (*ChAT::Cre; RphiGT*) to induce MN-specific G protein expression and enable monosynaptic labeling^49,50^. We injected RabiesΔG-mCherry unilaterally into the diaphragm of *Cdh9*^*GFP*^; *ChAT::Cre; RphiGT* mice at P4 to specifically label phrenic MNs and trace their synaptic inputs^47,51^ (Fig. 1B). Each injection labeled 1-9 starter phrenic MNs (Fig. S1A-B) which led to mCherry expression in 50-189 phrenic MN-targeting brainstem neurons (Fig. 1C, S1C). We quantified the overlap of GFP (induced in *Cdh9*^*GFP*^ mice) and mCherry (induced by rabies virus) expression in the brainstem at P11 (7 days after rabies injection) and found that 78.6 ± 2.5% of mCherry-labeled pre-motor neurons express GFP, indicating that the majority of phrenic MN-targeting rVRG neurons can be labeled in *Cdh9*^*GFP*^ mice (Fig. 1C-D).

While our tracing data indicated that *Cdh9*^*GFP*^ marks rVRG pre-motor neurons, *Cdh9* expression has also been reported in a subset of preBötC neurons^52^. Therefore, we assessed *Cdh9*^*GFP*^ expression throughout the VRC at P14 (Fig. 1E). We found that *Cdh9*^*GFP*^ labels the length of the rVRG, identified by its location dorsomedial to the loose nucleus ambiguus (NA) in the caudal ventrolateral medulla^21,45^. *Cdh9*^*GFP*^ expression rostral to the rVRG smoothly transitioned ventral to the semicompact NA, indicative of the preBötC^44,53^. Additionally, a small portion of these *Cdh9*^*GFP*^-positive neurons expressed SST, a marker of the preBötC^10-13^. Finally, *Cdh9*^*GFP*^ also appears to be expressed in the Bötzinger Complex (BötC), denoted by a further ventrolateral shift at the level of the compact NA^44^. Thus, our results show that *Cdh9*^*GFP*^ expression, in combination with local motor nuclei, can delineate multiple VRC populations, including phrenic MN-projecting rVRG neurons.

Since brainstem respiratory populations are heterogeneous, we next wanted to test whether *Cdh9*^*GFP*^ selectively targets either excitatory or inhibitory subpopulations. Excitatory neurons in both the preBötC and rVRG are derived from the ventral *Dbx1*-expressing p0 domain^10,11,20,21^ and can be targeted using a *Dbx1::Cre* driver^54^, while GABAergic inhibitory neurons can be labeled with *Vgat::Cre*^55^. We crossed *Cdh9*^*GFP*^ mice to mice expressing a Cre-dependent tdTomato reporter (*Ai9*)^56^ and either *Dbx1::Cre* (referred to as *Dbx1*^*tdTom*^) or *Vgat::Cre* (referred to as *Vgat*^*tdTom*^) to label both *Cdh9*-expressing excitatory or inhibitory neurons, respectively. We quantified the overlap of GFP and tdTomato in the rVRG at P0 and found that 73.1 ± 3.0% of Cdh9^GFP^ neurons overlap with the *Dbx1*-derived population (Fig. 1F-G), while 41.6 ± 4.2% are inhibitory (Fig. 1H-I). Our data indicate that *Cdh9*^*GFP*^-labeled neurons in the VRC encompass both inhibitory and excitatory populations, independent of their progenitor identity and function.

### Hox5 transcription factors delineate hindbrain respiratory populations

Since *Cdh9*^*GFP*^ mice provide a novel labeling tool for the VRC, we asked whether we could utilize these mice to identify transcriptional programs that selectively control the development and function of distinct respiratory populations. We confirmed that *Cdh9*^*GFP*^ mice show GFP expression in the VRC at embryonic timepoints, starting around e14.5-e15.5 (Fig. S2A). Since the VRC is organized rostrocaudally^1,2^, we asked whether Hox transcription factors, which are associated with early brainstem rostrocaudal patterning^57,58^, are differentially expressed in subsets of respiratory neurons. We initially tested the expression of Hox4 paralogs and Pbx co-factors, as they were previously reported to be enriched in *Dbx1*-derived brainstem neurons^39^. Although we observed robust expression of both Hoxc4 and Pbx1 in *Cdh9*^*GFP*^-labeled neurons at e15.5, these transcription factors were broadly expressed in both rostral preBötC and caudal rVRG neurons (Fig. S2B-D), suggesting they are unlikely to underlie the divergent development of these two populations.

Next, we assessed the expression of Hox5 proteins in the brainstem, as they have been previously reported to be expressed in the caudal medulla^40,41,59^. Since Hox5 proteins control the development of multiple respiratory neurons and tissues, including phrenic MNs, lungs, diaphragm, and trachea^60-65^, we hypothesized that they might also underlie aspects of brainstem respiratory neuron development. We observed Hoxa5 protein expression in the brainstem as early as e11.5 (data not shown). At e15.5, when respiratory populations can be demarcated in *Cdh9*^*GFP*^ mice, we found that Hoxa5, Hoxb5, and Hoxc5 are expressed in the majority of GFP+ rVRG neurons, while significantly fewer GFP+ preBötC neurons express any Hox5 paralog (Fig. 2A-F, K). This differential expression is also observed at birth (P0) and maintained at least through P14, although Hoxc5 cannot be detected in the medulla at postnatal stages (Fig 2G-J, L-M). Thus, maintained Hox5 expression delineates the rVRG from other medullary respiratory populations.

**Figure 2.**
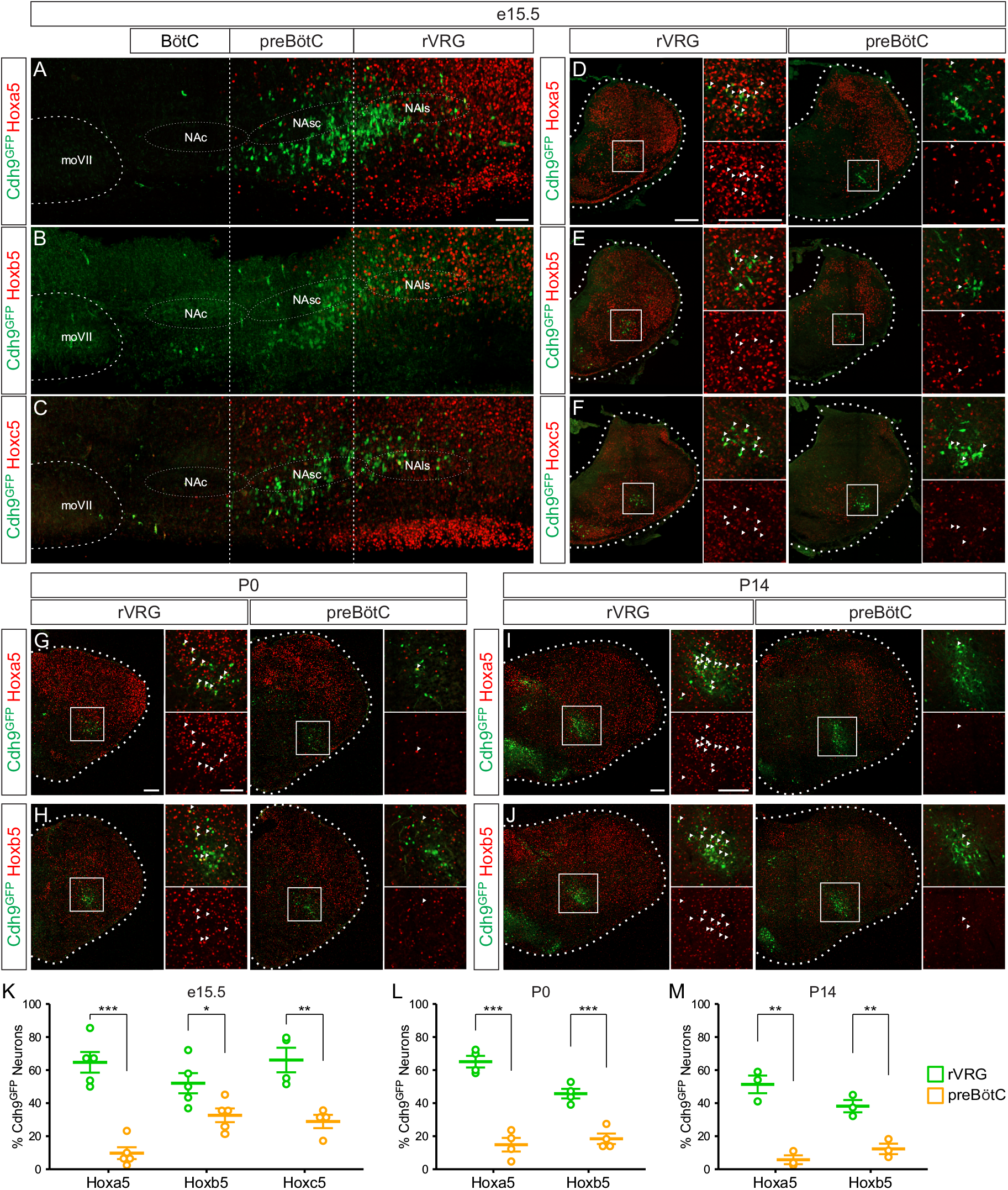
Hox5 expression delineates brainstem respiratory populations. A-C) Hoxa5 (A), Hoxb5 (B), and Hoxc5 (C) expression (red) in sagittal brainstem sections from *Cdh9*^*GFP*^ (green) mice at e15.5. Scale bar = 200 μm. D-F) Hoxa5 (D), Hoxb5 (E), and Hoxc5 (F) expression (red) in coronal brainstem sections from *Cdh9*^*GFP*^ (green) mice at e15.5. Transverse sections at the level of the rVRG and preBötC are shown. Scale bar = 200 μm. G-J) Hoxa5 (G, I) and Hoxb5 (H, J) expression (red) in coronal brainstem sections from *Cdh9*^*GFP*^ (green) mice at P0 (G-H) and P14 (I-J). Transverse sections at the level of the rVRG and preBötC are shown. Scale bar = 200 μm. (K) Percentage of rVRG and preBötC Cdh9^GFP^ neurons expressing Hox5 paralogs at e15.5 (Hoxa5 – rVRG: 64.7 ± 6.2%, preBötC: 9.7 ± 3.6%, n = 5, p = 0.0002; Hoxb5 – rVRG: 52.1 ± 6.1%, preBötC: 32.6 ± 4.2%, n = 5, p = 0.034; Hoxc5 – rVRG: 66.1 ± 7.4%, preBötC: 28.9 ± 4.0%, n = 4, p = 0.0084). (L) Percentage of rVRG and preBötC Cdh9^GFP^ neurons expressing Hox5 paralogs at P0 (Hoxa5 – rVRG: 65.1 ± 3.5%, preBötC: 14.8 ± 4.1%, n = 4, p = 0.0001; Hoxb5 – rVRG: 45.7 ± 2.9%, preBötC: 18.4 ± 3.2%, n = 4, p = 0.0007). (M) Percentage of rVRG and preBötC Cdh9^GFP^ neurons expressing Hox5 paralogs at P14 (Hoxa5 – rVRG: 51.4 ± 5.3%, preBötC: 5.7 ± 2.7%, n = 3, p = 0.005; Hoxb5 – rVRG: 38.2 ± 3.7%, preBötC: 12.3 ± 3.2%, n = 3, p = 0.0065).

### Selective deletion of *Hox5* genes in *Dbx1*-derived neurons leads to neonatal lethality

Given their selective expression in the rVRG and their critical role in phrenic MN development^60,63^, we asked whether Hox5 transcription factors are required in brainstem populations for respiratory circuit development and function. We used a *Dbx1::Cre* driver to selectively delete *Hoxa5*, the *Hox5* paralog most restricted to the rVRG, from excitatory respiratory neurons using a conditional *Hoxa5* allele (*Hoxa5*^*flox/flox*^; *Dbx1::Cre*)^54,66^. We introduced this conditional *Hoxa5* allele into a *Hoxb5*^*-/-*^ and *Hoxc5*^*-/-*^ background (*Hoxa5*^*flox/flox*^; *Hoxb5*^*-/-*^; *Hoxc5*^*-/-*^; *Dbx1::Cre*, referred to as *Hox5*^*Dbx1Δ*^) to eliminate all *Hox5* genes from excitatory brainstem respiratory neurons^67,68^. We validated our conditional knockout strategy at e15.5 via immunofluorescence for Hoxa5 and a Cre-dependent tdTomato reporter. In control embryos (*Hoxa5*^*flox/+*^; *Hoxb5*^*-/-*^; *Hoxc5*^*-/-*^; *Dbx1::Cre; Ai9*), we found 15.0 ± 1.6% overlap between Hoxa5 and tdTomato in the rVRG, which was nearly ablated in *Hox5*^*Dbx1Δ*^; *Ai9* embryos (Fig. S3A-B). We similarly confirmed the global deletion of *Hoxb5* and *Hoxc5* by immunofluorescence at e15.5 (Fig. S3C-D). Therefore, our approach successfully abolishes all *Hox5* expression in *Dbx1*-derived neurons. Though physically indiscernible from control littermates, neonatal (P0) *Hox5*^*Dbx1Δ*^ mice exhibit respiratory distress, with spasmic respiratory movements and visible activation of accessory muscles including the abdominals (Video S1). While mice that have at least a single *Hoxa5* or *Hoxb5* allele are viable and fertile, *Hox5*^*Dbx1Δ*^ mice die shortly after birth due to respiratory failure.

To ensure that neonatal lethality did not result from ectopic deletion of *Hox5* genes in phrenic MNs, we examined Hoxa5 expression in the spinal cord at e12.5. We found that the number of phrenic MNs expressing Hoxa5 was similar in control and *Hox5*^*Dbx1Δ*^ mice, which did not show a reduction in phrenic MN numbers (Fig. S4A-C). We also found that *Hox5*^*Dbx1Δ*^ mice show normal diaphragm innervation and no changes in the number of neuromuscular junctions (NMJs) at the diaphragm muscle at e18.5/P0 (Fig. S4D-E). Therefore, the neonatal lethality observed in *Hox5*^*Dbx1Δ*^ mice is not due to non-cell-autonomous effects on phrenic MN development, but rather due to the disruption of Hox5-dependent transcriptional programs in *Dbx1*-derived neurons.

### Robust respiration requires Hox5 activity in *Dbx1*-derived neurons

While *Hox5*^*Dbx1Δ*^ mice predominantly die shortly after birth, a small percentage survive until adulthood but appear to experience persistent respiratory distress. We measured the respiratory capacity of adult *Hox5*^*Dbx1Δ*^ mice and sex-matched, control littermates at P50 via unrestrained, whole body, flow through plethysmography (Fig. S5A-B)^63^. Under normal air conditions (79% N_2_, 21% O_2_), *Hox5*^*Dbx1Δ*^ mice exhibit a reduction in tidal volume (the amount of air inhaled during a normal breath) compared to their paired controls (Fig. S5C-D). However, an increase in respiratory frequency (Fig. S5E-F) enables *Hox5*^*Dbx1Δ*^ mice to maintain normal minute ventilation (volume of air inhaled per minute) at rest (Fig. S5G-H).

Despite maintaining breathing at rest, we observed that *Hox5*^*Dbx1Δ*^ mice tend to remain stationary, and if they are paired with a breeding partner, they perish by the next morning. Therefore, we hypothesized that *Hox5*^*Dbx1Δ*^ adult mice cannot maintain ventilation under heightened metabolic demands. To test this, we exposed *Hox5*^*Dbx1Δ*^ mice and their paired controls to a 5% CO_2_ hypercapnia challenge (74% N_2_, 21% O_2_, 5% CO_2_). In response to 5% CO_2_, control mice increase both their tidal volume (Fig. S5O) and their respiratory frequency (Fig. S5P) leading to a substantial increase in overall ventilation (Fig. S5Q). These responses are severely attenuated in *Hox5*^*Dbx1Δ*^ mice, which display only a modest increase in respiratory frequency and tidal volume (Fig. S5O-P), resulting in a small change in minute ventilation (Fig. S5Q). When compared to their littermate controls under hypercapnia, *Hox5*^*Dbx1Δ*^ mice exhibit a reduction in tidal volume (Fig. S5I-J), but without an accompanying increase in respiratory frequency (Fig. S5K-L), leading to an overall decrease in minute ventilation (Fig. S5M-N). Collectively, our plethysmography data indicate that Hox5 activity is required in *Dbx1*-derived respiratory populations to establish adequate breath depth. Under normal air conditions, compensatory mechanisms that enhance respiratory frequency can sometimes maintain ventilation to prevent neonatal death^69-74^; however, they are insufficient to sustain breathing under metabolic or atmospheric stress.

### Hox5 transcription factors are essential for patterned phrenic MN activation

As the diaphragm is the predominant inspiratory muscle, and phrenic MNs are the sole MN population responsible for diaphragm contractions, we hypothesized that impaired phrenic MN activation may underlie the reduced tidal volume in *Hox5*^*Dbx1Δ*^ mice. To determine whether Hox5-driven transcriptional programs are required in the core inspiratory circuit for precise phrenic MN activation, we obtained suction electrode recordings from the phrenic nerve of e18.5-P0 control and *Hox5*^*Dbx1Δ*^ mice using the reduced brainstem-spinal cord (*en bloc*) preparation (Fig. 3A)^63^. After removal of the pons to alleviate tonic inhibition onto the respiratory circuit, this preparation exhibits fictive breathing, defined as rhythmic bursts of phrenic nerve activity (Fig. 3B). Although burst frequency is highly variable in control preparations at this age, we observed a consistent decrease in *Hox5*^*Dbx1Δ*^ preparations, suggesting reduced respiratory frequency in *Hox5*^*Dbx1Δ*^ mice (Fig. 3C).

**Figure 3.**
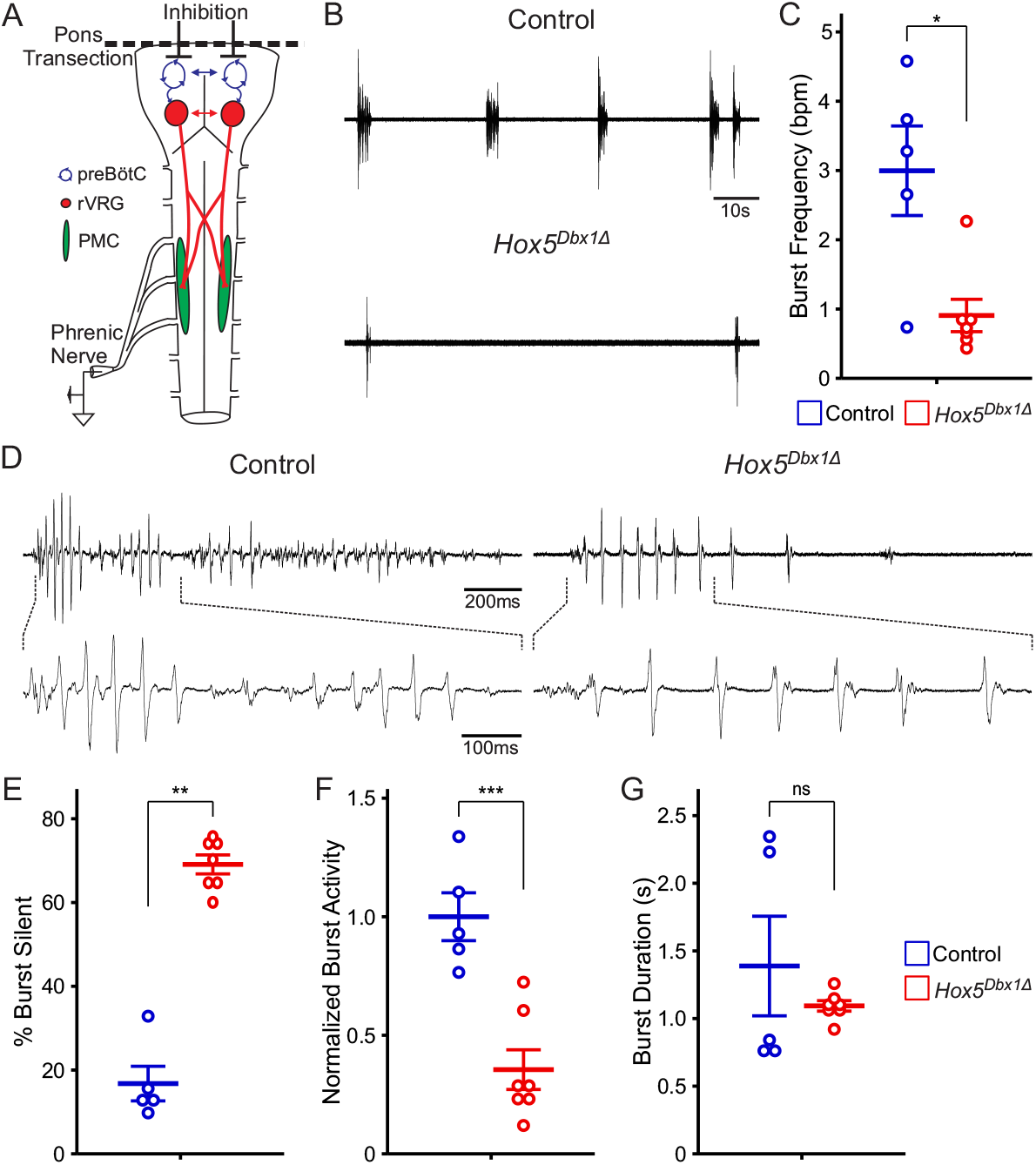
*Dbx1*-mediated *Hox5* deletion impairs phrenic nerve activity. (A) Schematic of the reduced medullary-spinal preparation used for phrenic nerve recordings. Transection of the pons alleviates tonic inhibition onto the respiratory circuit, inducing fictive breathing. Suction electrodes are attached to the phrenic nerve to record respiratory circuit output. (B) Traces of phrenic nerve activity from control and *Hox5*^*Dbx1Δ*^ preparations at P0. (C) Phrenic nerve burst frequency as bursts per minute (bpm) at P0 (Control: 3.00 ± 0.65, *Hox5*^*Dbx1Δ*^: 0.91 ± 0.23, n = 5-7, p = 0.018). (D) Single phrenic nerve bursts from control and *Hox5*^*Dbx1Δ*^ preparations at P0. Differences in burst patterning are highlighted below by expanding the denoted regions. (E) Mean quiescence within bursts at P0, shown as percent of burst with no unit activity (Control: 16.8 ± 4.1% of burst, *Hox5*^*Dbx1Δ*^: 69.1 ± 2.3% of burst, n = 5-7, p = 0.0025). (F) Mean phrenic nerve activity during bursts at P0, shown as integrated burst activity normalized to burst duration and to the average of controls (Control: 1.00 ± 0.10, *Hox5*^*Dbx1Δ*^: 0.36 ± 0.08, n = 5-7, p = 0.0009). (G) Mean burst duration at P0, shown in seconds (Control: 1.39 ± 0.37 seconds, *Hox5*^*Dbx1Δ*^: 1.09 ± 0.04 seconds, n = 5-7, p = 0.64).

We found that control bursts showed highly patterned MN activation, incorporating large spikes of synchronous motor unit activation and smaller, asynchronous peaks (Fig. 3D). In contrast, *Hox5*^*Dbx1Δ*^ bursts were far less complex, composed only of individual synchronous spikes separated by long periods of quiescence (Fig. 3D-E, S6A). These extended pauses coupled with shorter spike amplitude resulted in reduced phrenic MN activation, shown as integrated activity normalized to the average of control bursts (Fig. 3F, S6B). Despite changes in phrenic nerve burst frequency and composition, burst duration is unperturbed in *Hox5*^*Dbx1Δ*^ preparations (Fig. 3G, S6C). We noted that there is a bimodal distribution of control burst duration, however, burst duration remains consistent within preparations (Fig. S6C). We conclude that loss of Hox5 activity in *Dbx1*-derived neurons diminishes excitatory drive onto and disrupts coordinated activation of phrenic MNs.

### Loss of *Hox5* leads to a caudal expansion of SST+ putative preBötC neurons

What could be the cause of neonatal lethality and reduced phrenic MN activation in *Hox5*^*Dbx1Δ*^ mice? One potential explanation would be that *Hox5* deletion leads to a loss of brainstem respiratory neurons. To test this hypothesis, we performed immunofluorescence for Cleaved Caspase 3 (CC3), a member of the apoptotic cascade, in the developing brainstem to assess programmed cell death. We found no significant difference in CC3+ cells between control and *Hox5*^*Dbx1Δ*^ embryos at e11.5, the age at which Hox5 proteins are first detected in the brainstem (Fig. 4A-B). We also counted CC3+ cells at e18.5, when the respiratory circuit should be fully formed and shortly before *Hox5*^*Dbx1Δ*^ mice perish, and found no significant difference in apoptosis at this stage either (Fig. 4C-D), indicating that loss of *Hox5* does not result in the death of *Dbx1*-derived neurons in the brainstem.

**Figure 4.**
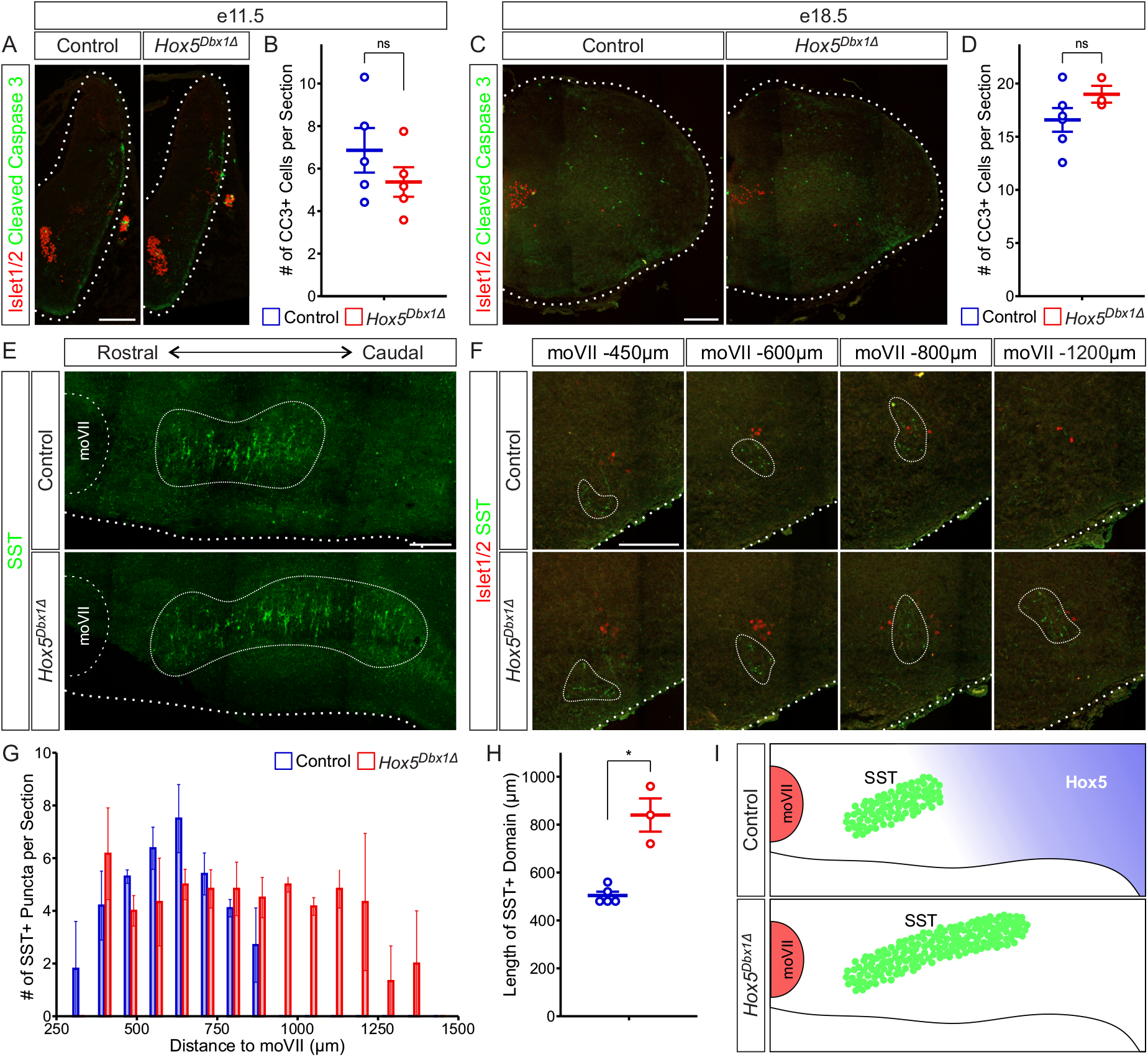
Hox5 transcription factors are critical for delineating VRC boundaries. A-D) Cleaved caspase 3 (CC3, green) immunofluorescence labels cells undergoing apoptosis in control and *Hox5*^*Dbx1Δ*^ embryos at e11.5 (A) and e18.5 (C). MNs are labeled by Islet1/2 (red). Scale bar = 200 μm. Average number of CC3+ cells per section at e11.5 (B, Control: 6.9 ± 1.0 cells, *Hox5*^*Dbx1Δ*^: 5.4 ± 0.7 cells, n = 5, p = 0.27) and e18.5 (D, Control: 16.6 ± 1.1 cells, *Hox5*^*Dbx1Δ*^: 19.0 ± 0.8 cells, n = 3-6, p = 0.12). (E) Sagittal brainstem sections in control and *Hox5*^*Dbx1Δ*^ mice at P0 showing caudal expansion of somatostatin (SST, green) expression in *Hox5*^*Dbx1Δ*^ mice at P0. Scale bar = 200 μm. (F) Coronal sections through the VRC of control and *Hox5*^*Dbx1Δ*^ mice at P0 showing SST (green) localization with respect to the NA (Islet1/2, red). The rostrocaudal position of each section was determined by distance to moVII. Scale bar = 200 μm. (G) Distribution of SST+ neurons throughout the VRC of control and *Hox5*^*Dbx1Δ*^ mice at P0 by distance to moVII (μm) (n = 3-5). (H) Rostrocaudal length of the SST+ domain in the VRC of control and *Hox5*^*Dbx1Δ*^ mice at P0 (Control: 504 ± 16 μm, *Hox5*^*Dbx1Δ*^: 840 ± 69 μm, n = 3-5, p = 0.032). (I) Schematic demonstrating the relationship between Hox5 expression and SST+ neurons. In control animals, the rostral boundary of Hox5 delineates the caudal boundary of SST expression in the VRC. In *Hox5*^*Dbx1Δ*^ mice, this boundary is disrupted, leading to a caudal expansion of the SST+ domain.

*Hox* gene deletion in spinal MNs often leads to a switch in MN subtype identity and ectopic MN expansion^57,75,76^, analogous to homeotic transformations of *Hox* mutants along the body axis^77^. Given the changes in both the integrated phrenic MN activity and burst frequency in our suction electrode recordings, we asked whether there might be a switch in the identity of brainstem respiratory neurons leading to changes in their proportion and distribution. Since *Hox5* expression delineates the boundary between the rVRG and preBötC, we hypothesized that this boundary may be eroded in *Hox5*^*Dbx1Δ*^ mice. To test this, we performed immunofluorescence for SST, which labels a subset of preBötC neurons^10-13^, in sagittal sections of control and *Hox5*^*Dbx1Δ*^ brainstems. While SST expression is highly restricted to the preBötC in controls, we found a dramatic elongation of the SST-expressing domain in *Hox5*^*Dbx1Δ*^ mice at P0 (Fig. 4E).

To quantify the expansion of SST+ neurons, we obtained coronal sections throughout the brainstem, denoted by their distance caudal to the facial motor nucleus (moVII), at P0 (Fig. 4F). In control mice, SST+ neurons can be detected up to 800 μm caudal to moVII, yet they are present as far as 1200 μm from moVII in *Hox5*^*Dbx1Δ*^ mice (Fig. 4G). We found that the length of the SST+ region is 504 ± 16 μm in control mice but expands to 840 ± 69 μm in *Hox5*^*Dbx1Δ*^ mice (Fig. 4H). To determine the directionality of this expansion, we measured the distance from moVII to the most rostral SST+ neurons and from the most caudal SST+ neurons to the spinal cord. In control mice, SST+ neurons are located 616 ± 20 μm away from the spinal cord, while SST-expressing neurons extend within 240 ± 69 μm of the spinal cord in *Hox5*^*Dbx1Δ*^ mice (Fig. S7A). On the contrary, the distance between the most rostral SST+ neurons and moVII remains consistent in control and *Hox5*^*Dbx1Δ*^ mice (Fig. S7B). Altogether, our data show that conditional deletion of *Hox5* from *Dbx1*-derived neurons leads to a caudal expansion of SST+ putative preBötC neurons, likely at the expense of the rVRG, and a degradation of the boundary delineating the rVRG and preBötC (Fig. 4I).

## Discussion

Brainstem respiratory neurons must be generated at the correct position and numbers and become highly specialized to support the critical function of breathing. Here, we find that Hox5 transcription factor activity is required in *Dbx1*-derived neurons in a cell-autonomous manner for proper respiratory circuit development and function.

### Cdh9 is a broad molecular marker for VRC neurons

While it has been reported that Cdh9+ neurons are a specialized subset of preBötC neurons involved in arousal^52^, we find that the *Cdh9*^*GFP*^ reporter mouse labels populations of VRC neurons broadly, including phrenic-projecting rVRG and SST+ preBötC neurons. Our findings may reflect temporally distinct phases of *Cdh9* expression, where *Cdh9* is initially expressed throughout the VRC while later being restricted to arousal-related preBötC neurons. Both Cdh9 function and the regulatory mechanisms that induce *Cdh9* expression in respiratory neurons are unclear. We find that *Cdh9* is expressed in both excitatory, *Dbx1*-derived, and inhibitory, likely *Lbx1*-derived^21^, rVRG neurons, as well as in the Hox5+ rVRG and Hox5-preBötC, making it unlikely that progenitor-restricted or Hox-dependent programs alone induce its expression. Rather, *Cdh9* expression may be regulated by the intersection of multiple transcriptional programs. Functionally, Cdh9 may enable selective connectivity between preBötC and rVRG neurons, or more broadly throughout the VRC, expanding its previously described role in rVRG-phrenic MN connectivity^47^. This function would likely require combinatorial expression and interaction with other cadherins, as we previously found that ablation of all cadherin signaling from *Dbx1*-derived neurons leads to perinatal lethality, but *Cdh9* knockout mice exhibit largely normal breathing behaviors^47^.

### Functional specialization of excitatory VRC populations

Dbx1+ neural progenitors give rise to at least two functionally distinct classes of excitatory neurons in the preBötC, as well as excitatory rVRG neurons, all of which are essential for respiration^10,11,20,21^. Type 1 preBötC neurons fire spontaneously, as recurrent connections lead to synchronous pre-inspiratory burstlets and recruitment of SST+ type 2 neurons^6^. Type 2 SST+ neurons send afferents to pre-motor neurons within the rVRG, with sufficient activation leading to inspiratory bursts and MN activation^6^. Our data show that loss of *Hox5* transcription factors in *Dbx1*-derived neurons causes a caudal expansion of SST+ neurons. It is unclear how these ectopic neurons are integrated into the respiratory circuit and whether there is also a concurrent increase in Type 1 neurons. Surprisingly, we find that an expansion in putative preBötC neurons leads to a reduction in phrenic nerve burst frequency. This might be due to reduced recurrent connectivity among preBötC neurons, requiring the activation of more neurons to drive full inspiratory bursts. Alternatively, the increase in excitatory preBötC compared to its unperturbed inhibitory component could lead to hyperexcitability, resulting in prolonged refractory periods and reduced inspiratory burst frequency^70,78,79^. Interestingly, there is a significant increase in respiratory frequency in *in vivo* plethysmography recordings, likely due to the presence of sensory feedback and chemosensory modulation which can compensate for changes in tidal volume in the intact animal^69-74^.

Regardless of whether only type 2 or all excitatory preBötC neurons expand caudally in *Hox5*^*Dbx1Δ*^ mice, this expansion likely comes at the expense of *Dbx1*-derived rVRG neurons, leading to reduced excitatory drive onto phrenic MNs. This is consistent with both reduced tidal volume *in vivo* and reduced phrenic MN activation *ex vivo*. Loss of excitatory rVRG neurons also provides an explanation for the altered phrenic nerve burst composition seen in *Hox5*^*Dbx1Δ*^ preparations. Balanced excitatory and inhibitory input onto phrenic MNs is critical for coordinated MN activation and efficient diaphragm contractions, as blockade of spinal GABAergic and glycinergic receptors results in a loss of synchronous MN activation^63^. We find that loss of Hox5 activity in *Dbx1*-derived rVRG neurons causes excessive synchronization of phrenic MN activity, suggesting a shift in the excitatory-inhibitory balance of phrenic MN inputs, leading to reduced and re-patterned phrenic MN activation.

### *Hox5* genes in the development of respiratory circuits

Early expression of Hox proteins in the developing brainstem establishes hindbrain segmentation and is essential for the rostrocaudal organization and proper function of respiratory circuits^57^. Hox1 and Hox2 paralogs are required for the development of rostral respiratory populations such as the pFRG and pontine respiratory neurons, respectively, but their deletion also leads to broad defects in rhombomere segregation and brainstem development^33-37^. Here, we find that *Hox5* deletion in *Dbx1*-derived neurons also leads to perinatal lethality due to respiratory failure. However, the temporal requirement for Hox5 appears to be delayed compared to other brainstem Hox proteins, as Hoxa5 protein is not detectable until e11.5, after rhombomere segregation has been completed, and is maintained through postnatal stages. Thus, it is likely that Hox5-driven transcriptional programs are acting in post-mitotic neurons to specify distinct neuronal subtype identities, analogous to Hox protein function in spinal MN development.

Hox5 transcription factors are also essential for the development and specification of phrenic MNs in the cervical spinal cord^60,63,64^. While *Hoxb5* is not expressed in MNs at this level, deletion of *Hoxa5* and *Hoxc5* eliminates nearly all phrenic MNs, resulting in persistent apnea and perinatal death^60^. Transcriptomic analyses showed that Hox5 proteins regulate molecular pathways essential for the unique dendrite arborization, axon pathfinding, and synaptic connectivity that differentiate phrenic MNs from all other MNs^63,64^. The sustained expression of Hox5 proteins in the rVRG suggests a similar role in establishing rVRG-specific properties, such as long-range bulbospinal projections to phrenic MNs, although the specific processes and molecular pathways under Hox5 control remain to be determined. In addition to specifying rVRG properties, alternative programs that define other respiratory populations, for example SST expression in the preBötC, must be repressed. In the spinal cord, cross-repression among different Hox paralogs ensures sharp boundaries between MN subtypes^75,76^. For example, Hoxc9 suppresses the expression of more rostral *Hox* genes to establish thoracic MN identity^80^. However, this does not seem to be the case for brainstem respiratory neurons, as there is not a sharp boundary between Hox4 and Hox5 which show largely overlapping expression in rVRG neurons. While Hox5 proteins do not appear to repress rostral *Hox4* paralog genes, they may function with yet unidentified cofactors to repress preBötC identity. In the cervical spinal cord, Hoxa5 forms a complex with Pbx cofactors and either Scip or FoxP1 to induce phrenic or limb MN identity, respectively, and repress the alternative fate^64^. While Pbx proteins are broadly expressed throughout the brainstem, we did not detect Scip expression in the VRC at early postnatal stages (data not shown) indicating that distinct Hox5 cofactors underlie Dbx1+ interneuron and MN specification. In addition to *Dbx1*-derived brainstem respiratory neurons and phrenic MNs, Hox5 proteins control the development of many structures throughout the respiratory system, including the diaphragm, lungs and trachea, indicating a central role for Hox5 as a master regulator for the respiratory system^61,62,65^.

### STAR Methods

#### Mouse Genetics

*Cdh9::FlpO* mice were generated using CRISPR-Cas9 technology. Briefly, a *P2A-FlpO* sequence (1468 bp) was inserted in place of the stop codon in mCdh9 exon 12. sgRNAs targeting within 20 bp of the desired integration site were designed with at least 3 bp of mismatch between the target site and any other site in the genome. Targeted integration was confirmed in vitro prior to moving forward with embryo injections. C57BL/6J fertilized zygotes were micro-injected in the pronucleus with a mixture of Cas9 protein at 30-60 ng/μl, and single guide RNA at 10-20 ng/μl each, and a ssDNA at 5-10 ng/μl (a ribonucleoprotein complex). The injected zygotes, after culture in M16 or alternatively Advanced-KSOM media, were transferred into the oviducts of pseudo-pregnant CD-1 females. Founder mice were genotyped by targeted next generation sequencing followed by analysis using CRIS.py. The following lines were generated as previously described and maintained on a mixed background: *RphiGT* (JAX# 024708)^49^, *ROSA26Sor*^*tm9(CAG-tdTomato)*^ (Ai9, JAX# 007909)^56^, *ChAT::Cre*^*50*^, *Dbx1::Cre*^54^, *RCE:FRT* (MMRRC# 032038-JAX)^48^, *Vgat::Cre* (JAX# 016962)^55^, *loxP*-flanked *Hoxa5* (*Hoxa5*^*flox/flox*^)^*66*^, *Hoxb5-/-*^67^, *Hoxc5-/-*^*68*^. Mouse colony maintenance and handling strategies adhered to protocols approved by the Institutional Animal Care Use Committee (IACUC) of Case Western Reserve University (assurance number: A-3145-01, protocol number: 2015-0180). Mice were housed in microisolator cages under a 12-hour light/dark cycle. Each cage contained no more than 2 adult mice with 1 litter or 5 adult mice.

#### Tissue Preparation

Neonatal (P0) mice were dissected acutely after cryo-anesthetization. Mice older than P10 were anesthetized via an intraperitoneal injection of a ketamine/xylazine cocktail. P14 mice were additionally perfused with ice-cold phosphate buffered saline (PBS) followed by ice-cold 4% paraformaldehyde (PFA). All postnatal brainstems and spinal cords were fixed in 4% PFA overnight at 4 °C. For embryonic tissue, pregnant mice (determined by vaginal plug) were euthanized via CO_2_ asphyxiation. Embryos (e11.5-e18.5) were dissected in ice-cold PBS and fixed in 4% PFA for 2 hours at 4 °C. Perinatal (e18.5-P0) diaphragms were pinned flat and fixed in 1% PFA for 1 hour at room temperature. All tissue was washed in PBS following fixation. For sectioning on a Leica VT1000S vibratome, fixed tissue was embedded in 4% low melting point agarose (Invitrogen). Consecutive 100 μm sections were taken and stored in PBS containing 0.02% sodium azide. For sectioning with a CM3050S Leica cryostat, tissue was dehydrated in 30% sucrose overnight, embedded in Optimal Cutting Temperature (OCT) compound, and stored at -80 °C. Transverse 16 μm cryosections of the brainstem and/or spinal cord were mounted serially across 5 replicate Superfrost plus gold glass slides (Thermo Fisher Scientific).

#### Rabies Virus Production and Monosynaptic Tracing

RabiesΔG-mCherry virus production and monosynaptic tracing were performed as previously described^47,81^ Rabies injection solution was made by mixing RabiesΔG-mCherry virus (titer of around 1^10^ TU/ml) with silk fibroin (Sigma-Aldrich# 5154)^82^ at a 2:1 ratio. 1-1.5 μl of rabies injection solution was unilaterally injected into the diaphragm of P4 cryo-anesthetized *ChAT::Cre; RphiGT*; *Cdh9::FlpO; RCE:FRT* mice using a nano-injector (Drummond). Mice were sacrificed 7 days post-injection (P11), and brainstem sections were obtained using a vibratome. Specific labeling of ipsilateral phrenic MNs was confirmed by monitoring mCherry fluorescence in the diaphragm and spinal cord. Samples with phrenic MN-specific labeling only showed fluorescent signal at C3-C5 ventral roots.

#### Immunohistochemistry

Immunohistochemistry was performed as previously described^60,64^. Cryo or vibratome sections were permeabilized and blocked with PBS containing Triton X-100 (0.1% PBT for 16 μm/0.5% PBT for 100 μm) and 1% bovine serum albumin (BSA). Sections were incubated with primary antibodies diluted in PBT and 0.1% bovine serum albumin (BSA) overnight (4 °C/room temperature), washed 3 times with PBT, and incubated with secondary antibodies in PBT (1 hour/overnight) at room temperature followed by 3 more washes of PBT. After a final wash in PBS, vibratome sections were mounted sequentially onto Superfrost plus gold glass slides (Thermo Scientific). Vectashield Vibrance mounting medium (Vector Laboratories) and cover glass (VWR) were applied to preserve fluorescence.

Diaphragm whole mounts were stained as described^63^. Following fixation, the tissue was permeabilized and blocked in 0.5% PBT and 4% normal goat serum (blocking solution) for 1 hour at room temperature. Primary antibodies were diluted in blocking solution and applied overnight at 4 °C. Tissue was washed 3 times with 0.5% PBT, then incubated with secondary antibodies diluted in blocking solution overnight at 4 °C. Finally, diaphragms were washed 3 times with 0.5% PBT, once with PBS, then briefly post-fixed with 1% PFA before mounting and cover slipping.

The following primary antibodies were used in this study: mouse anti-Islet1/2 (1:1000; DSHB, RRID: AB_2314683)^83^, goat anti-ChAT (1:250; MilliporeSigma, RRID: AB_2079751), rabbit anti-SST (1:300; BMA Biomedicals, RRID: AB_518614), rabbit anti-GFP (1:1000; Thermo Fisher Scientific, RRID: AB_221570), rabbit anti-DsRed (1:1000; Takara Bio, RRID: AB_10013483), sheep anti-digoxigenin-POD fab fragments (1:3000, Roche, cat# 11207733910), guinea pig anti-Hoxa5 (1:1000)^76^, rabbit anti-Hoxb5 (1:1000)^60^, guinea pig anti-Hoxc5 (1:1000)^76^, rabbit anti-Hoxc4 (1:1000)^76^, rabbit anti-Pbx1 (1:1000; Cell Signaling Technology, RRID:AB_2160295), goat anti-Scip (1:5000; Santa Cruz Biotechnology, RRID: AB_2268536), α-bungarotoxin Alexa Fluor 555 conjugate (1:1000; Thermo Fisher Scientific, RRID: AB_2617152), rabbit anti-neurofilament (1:1000; Synaptic Systems, RRID: AB_887743), and rabbit anti-cleaved caspase 3 (1:1000; Cell Signaling, RRID: AB_2341188).

The following fluorophore-conjugated secondary antibodies were used in this study: donkey anti-goat Alexa Fluor 488 (Thermo Fisher Scientific, RRID: AB_2534102), donkey anti-goat Alexa Fluor 647 (Jackson ImmunoResearch Labs, RRID: AB_2340437), donkey anti-rabbit Alexa Fluor 488 (Abcam, RRID: AB_2636877), donkey anti-rabbit Alexa Fluor 555 (Thermo Fisher Scientific, RRID: AB_162543), donkey anti-rabbit Alexa Fluor 647 (Thermo Fisher Scientific, RRID: AB_2536183), donkey anti-guinea pig CY3 (Jackson ImmunoResearch Labs, RRID: AB_2340460) and donkey anti-guinea pig Alexa Fluor 647 (Jackson ImmunoResearch Labs, RRID: AB_2340476).

#### Confocal Microscopy and Image Processing

Fluorescent images were captured with a Zeiss LSM 800 laser scanning confocal microscope equipped with a 20x/0.8 apochromatic objective lens and a 40x/1.3 oil apochromatic objective lens using the 405, 488, 555/561, and 647 nm laser lines. To capture the full extent of fluorescence within the target tissue, serial Z-stacks were captured through a tiled region of interest. Data were processed using Zen (blue edition) software to stitch tiled images and convert Z-stacks into a 2D representation via maximum intensity orthogonal projection.

Overlap of Cdh9^GFP^ neurons with Ai9 or immunofluorescent signal was quantified visually in Zen (blue edition) software. Overlap of *Dbx1::Cre*-dependent tdTomato with Hoxa5 was quantified using the JACoP plugin for Fiji^84^ as the Mander’s Coefficient with automatic thresholding. Phrenic MNs were identified bilaterally based on the co-expression of Islet1/2 and Scip, with Hoxa5 signal being manually assessed. Diaphragm Innervation was quantified using Volocity (RRID: SCR_002668) to detect α-bungarotoxin signal, apply a fine noise filter, and count objects larger than 25 μm^2^.

#### Definition of VRC Populations

Ventral respiratory column (VRC) neurons have been delineated in postnatal tissue via retrograde tracing and electrophysiological studies^44-46^. Based on these studies, we found that the Bötzinger Complex (BötC), pre-Bötzinger Complex (preBötC), and rostral Ventral Respiratory Group (rVRG) were all labelled and could be distinguished in *Cdh9*^*GFP*^ reporter mice in tandem with local MN populations at P14. Thus, we defined the rVRG as Cdh9^GFP^ neurons located medial to the loose Nucleus Ambiguus (NA). The preBötC was defined just rostral to the rVRG, where Cdh9^GFP^ neurons shift ventral to the semicompact NA forming an elongated dorsoventral strip. Finally, the BötC was defined as further rostral Cdh9^GFP^ neurons located ventral and distal to the compact NA. These topographical definitions were applied to embryonic stages, where classical definitions based on distance to the facial motor nucleus (moVII) are insufficient, as brainstem growth during development renders measurement-based definitions unreliable.

In experiments which did not include the *Cdh9*^*GFP*^ reporter, we defined the rVRG by distance to the spinal cord using data from age-matched *Cdh9*^*GFP*^ reporter experiments. These definitions were corroborated by using Hoxa5 expression as an additional landmark, which was most critical for experiments at e11.5, which is prior to the organization of the NA and expression of *Cdh9*^*GFP*^. At e11.5, we defined our region of interest to be within 480 μm (6 replicate sections) of the spinal cord. At e15.5, we defined the rVRG as being within 480 μm (6 replicate sections) of the spinal cord. At e18.5, we defined the rVRG as being within 720 μm (9 replicate sections) of the spinal cord.

#### Whole body plethysmography

Plethysmography was performed on conscious, unrestrained adult mice (P50) as previously described^51,63^. One *Hox5*^*Dbx1Δ*^ mouse and one control (*Hoxa5*^*flox/flox*^; *Hoxb5*^*+/-*^; *Hoxc5*^*-/-*^; *Dbx1::Cre*), sex-matched littermate were simultaneously placed into individual whole body, flow through plethysmograph chambers (emka) attached to differential pressure transducers (emka). Chambers were filled with “normal air” (79% N_2_, 21% O_2_) at a flow rate of 0.75 L/minute per chamber. Mice were allowed to acclimate until calm (∼1-2 hours), at which point the provided air was changed to “5% CO_2_” (74% N_2_, 21% O_2_, 5% CO_2_). This hypercapnic condition was maintained for 30 minutes. Breathing parameters including frequency, tidal volume, and minute ventilation were recorded using the iox2 software (emka). This paradigm was repeated on 3 consecutive days, rotating the chambers used to prevent technical bias.

All breaths were collected and visualized over time. As respiration can greatly vary based on the subject’s activity, periods of resting (characterized by a consistent pattern of low frequency breaths) were manually selected across all trials. Additionally, breaths occurring within 10 minutes of hypercapnic exposure were omitted, as gas exchange within the chamber is a gradual process. All qualified breaths were used to determine frequency, tidal volume, and minute ventilation distribution per animal. Mean respiratory values in each atmospheric condition were normalized to that of the paired littermate control, shown as percent of control. Additionally, mean values from 5% CO_2_ were normalized to their values in normal air within sample, shown as percent change.

#### Electrophysiology

Suction electrode recordings from the phrenic nerve were performed as previously described^63,85^. Neonatal mice were cryoanesthetized and rapidly dissected in oxygenated Ringer’s solution (128 mM NaCl, 4 mM KCl, 21 mM NaHCO_3_, 0.5 mM NaH_2_PO_4_, 2 mM CaCl_2_, 1 mM MgCl_2_, and 30 mM D-glucose; equilibrated by bubbling in 95% O_2_/5% CO_2_). The brainstem and cervical spinal cord were exposed via ventral laminectomy, and the phrenic nerves were dissected free of connective tissue. Removing the pons by cutting at the pontomedullary boundary initiated fictive inspiration, evident by intercostal and diaphragm contractions. The phrenic nerves were clipped proximal to the diaphragm and pulled into suction electrodes. Recordings were performed with continuous perfusion of oxygenated Ringer’s solution. Signal was band-pass filtered from 10 Hz to 3 kHz using Grass amplifiers, amplified 5,000-fold, and sampled at a rate of 50 kHz with a Digidata 1440A (Molecular Devices). The resulting traces were visualized and recorded using Axoscope (Molecular Devices). Spike2 (Cambridge Electronic Design) was used to analyze burst duration, burst frequency, integrated burst activity, and percent of burst with no activity. These parameters were assessed for 5-7 bursts per preparation which were averaged to represent the phrenic nerve activity of a single sample.

#### SST Quantification

Quantification of somatostatin (SST) localization was performed in serial coronal sections of the perinatal brainstem. Islet1/2 and choline acetyltransferase (ChAT) were used to visualize local motor neurons including moVII and NA. The number of SST puncta per section were counted visually in Zen (blue edition) software with reference to the ipsilateral moVII. The number of sections containing clear SST puncta were counted to determine the length of the SST+ domain. To calculate the distance of the SST+ domain to the spinal cord, the number of sections between the caudal-most SST+ and rostral-most spinal cord sections was counted. The number of sections between the rostral-most SST+ and caudal-most moVII sections was used to determine the distance of the SST+ domain to moVII.

#### Experimental Design and Statistical Analysis

To maintain rigor and reproducibility, male and female mice were used interchangeably for all experiments. Genotypes for control mice in conditional knockout experiments include *Hoxa5*^*flox/flox*^; *Hoxb5*^*-/-*^; *Hoxc5*^*-/-*^ and *Hoxa5*^*flox/delta*^; *Hoxb5*^*-/-*^; *Hoxc5*^*-/-*^, *Hoxa5*^*flox/flox*^; *Hoxb5*^*+/-*^; *Hoxc5*^*-/-*^ and *Hoxa5*^*flox/delta*^; *Hoxb5*^*+/-*^; *Hoxc5*^*-/-*^. Genotypes for experimental (*Hox5*^*Dbx1Δ*^) mice in conditional knockout experiments include *Hoxa5*^*flox/flox*^; *Hoxb5*^*-/-*^; *Hoxc5*^*-/-*^; *Dbx1::Cre* and *Hoxa5*^*flox/delta*^; *Hoxb5*^*-/-*^; *Hoxc5*^*-/-*^; *Dbx1::Cre*. We sometimes noted germline recombination when breeding *Dbx1::Cre* mice, leading to pups with a *Hoxa5*^*delta*^ allele. We found *Hoxa5*^*delta/flox*^ mice to be indistinguishable from *Hoxa5*^*flox/flox*^ littermates. Data were presented as box or line plots with each point representing data from an individual mouse unless otherwise noted. As all experiments compared only two groups, Shapiro Wilk tests were used to determine normality. For normally distributed data, unpaired Student’s t-tests were performed. One sample Student’s t-tests (hypothetical value = 0) were performed for data which was normalized to paired control samples. For data not distributed normally, the Wilcoxon Rank-Sum test was used instead. Data are represented as the mean ± standard error. Statistical analyses and plot generation were performed in R (v4.1.1). Statistical significance was achieved with p < 0.05, where *p < 0.05, **p < 0.01, ***p < 0.001, and ****p < 0.0001.

### Resource availability

#### Lead contact

Requests for further information and resources should be directed to the lead contact, Polyxeni Philippidou.

#### Materials availability

*Cdh9::FlpO* mice were generated by Lindsay A. Schwarz and are available upon request.

#### Data and code availability

The data generated in this study are available from the lead contact upon request. The code used for plethysmography analysis will be available at GitHub and will be publicly available as of the date of publication. Any additional information required to reanalyze the data reported in this paper is available from the lead contact upon request.

## Acknowledgements

We thank Deneen Wellik for providing *Hoxb5-/-*mice and Jeremy Dasen for Hox5 antibodies. This work was funded by NIH R01NS114510 to PP, institutional funds from St. Jude Children’s Research Hospital to LAS, F31NS124240 to MTM, F30HD096788 to ANV, T32GM007250 to ANV/CWRU MSTP. PP is the Weidenthal Family Designated Professor in Career Development.

## Author contributions

MTM and PP conceived and designed the study, MTM, ML, ANV, RLB and EMB performed experiments, MTM, ML, ANV, EMB and PP analyzed data, LJ provided Hoxa5 floxed mice, LAS generated *Cdh9::FlpO* mice, and MTM and PP wrote the paper with input from all authors.

## Supplemental information

Document S1. Figures S1–S7

Video S1. Breathing behavior of a *Hox5*^*Dbx1Δ*^ pup and littermate control at P0, related to Figure 3

