## Supplemental Figures for "A sustained Hox program delineates brainstem neurons essential for breathing"

##### Figure S1. Monosynaptic retrograde rabies virus tracing of phrenic MNs

A) Example of a starter phrenic MN expressing mCherry (red) and choline acetyltransferase (ChAT, green) at P11. Scale bar = 200 μm.

B) Quantification of starter (mCherry+) phrenic MNs at P11 (3.4 ± 1.4 labeled phrenic MNs, n = 5).

C) Quantification of mCherry+ neurons in the brainstem at P11 (92.2 ± 25.7 mCherry+ neurons, n = 5).

##### Figure S2. Hoxc4 and Pbx1 are broadly expressed in the rVRG and preBötC

A) Consecutive sagittal sections (top = medial, bottom = lateral) through the VRC of a $Cdh9^{GFP}$ (green) mouse at e15.5. Islet1/2 (red) labels the facial motor nucleus (moVII) and the nucleus ambiguus (NA). Scale bar = 200 μm

B) Brainstem Hoxc4 (top) and Pbx1 (bottom) (red) expression in $Cdh9^{GFP}$ (green) mice at e15.5. Transverse sections at the level of the rVRG and preBötC are shown. Scale bar = 200 μm.

C) Quantification of Hoxc4 expression in $Cdh9^{GFP}$ neurons at e15.5 (rVRG: 79.7 ± 3.1% of $Cdh9^{GFP}$ neurons, preBötC: 88.2 ± 2.2% of $Cdh9^{GFP}$ neurons, n = 4, p = 0.072).

D) Quantification of Pbx1 expression in $Cdh9^{GFP}$ neurons at e15.5 (rVRG: 69.8 ± 4.1% of $Cdh9^{GFP}$ neurons, preBötC: 71.3 ± 5.2% of $Cdh9^{GFP}$ neurons, n = 4, p = 0.88).

##### Figure S3. Validation of $Hox5^{Dbx1\Delta}$ mice

A) Hoxa5 (green) expression in control and $Hox5^{Dbx1\Delta}$ brainstem sections at the level of the rVRG at e15.5. *Dbx1::Cre*-mediated tdTomato ($Dbx1^{tdTom}$, red) marks Dbx1-derived neurons. Scale bar = 100 μm.

B) Percentage of $Dbx1^{tdTom}$ pixels overlapping with Hoxa5 at e15.5 (Control: 15.0% ± 1.6%, $Hox5^{Dbx1\Delta}$: 0.9% ± 0.1%, n = 2-3, p = 0.013).

C-D) Hoxb5 (C) and Hoxc5 (D) (green) expression in control and $Hox5^{Dbx1\Delta}$ brainstem sections at the level of the rVRG at e15.5. Islet1/2 (red) labels MNs. Scale bar = 100 μm.

##### Figure S4. Normal phrenic MN development in $Hox5^{Dbx1\Delta}$ mice

A) Hoxa5 (green) expression in phrenic MNs, identified by Scip (red), in control and $Hox5^{Dbx1\Delta}$ embryos at e12.5. Scale bar = 100 μm.

B) Percentage of phrenic MNs expressing Hoxa5 at e12.5 (Control: 93.2% ± 1.2%, $Hox5^{Dbx1\Delta}$: 92.1% ± 1.7%, n = 3, p = 0.95).

C) Quantification of bilateral phrenic MNs at e12.5 (Control: 1640.8 ± 174.8 phrenic MNs, $Hox5^{Dbx1\Delta}$: 1656.7 ± 175.4 phrenic MNs, n = 3, p = 0.62).

D) Diaphragm innervation in control and *Hox5*$^{Dbx1Δ}$ mice at e18.5. Phrenic motor axons are labeled by neurofilament (green) and neuromuscular junctions (NMJs) by α-bungarotoxin (red). Scale bar = 250 μm.

E) Quantification of NMJs in control and *Hox5*$^{Dbx1Δ}$ diaphragms (Control: 6036.6 ± 592.6 NMJs, *Hox5*$^{Dbx1Δ}$: 4472.3 ± 595.8 NMJs, n = 6-8, p = 0.88)

**Figure S5. Hox5 activity is required in Dbx1-derived neurons for robust and adaptable respiration**

A) Schematic of unrestrained, whole body, flow-through plethysmography. A *Hox5*$^{Dbx1Δ}$ mouse and a sex-matched control littermate are placed in airtight chambers to detect changes in air pressure due to respiration. Mice are provided normal air for 1 hour followed by 5% $CO_2$ for 20 minutes while resting.

B) Plethysmography traces from control (left) and *Hox5*$^{Dbx1Δ}$ (right) mice in normal air (top) or hypercapnia (bottom) at P50.

C) Distribution plot of breath tidal volume for control and *Hox5*$^{Dbx1Δ}$ mice in normal air at P50 (n = 5, ribbon = standard error).

D) Mean tidal volume, shown as percent of sex-matched control littermates in normal air at P50 (*Hox5*$^{Dbx1Δ}$: 26.6 ± 6.0% decrease, n = 5, p = 0.011).

E) Distribution plot of breath frequency for control and *Hox5*$^{Dbx1Δ}$ mice in normal air at P50 (n = 5, ribbon = standard error).

F) Mean breath frequency, shown as percent of sex-matched control littermates in normal air at P50 (*Hox5*$^{Dbx1Δ}$: 52.1 ± 10.2% increase, n = 5, p = 0.007).

G) Distribution plot of breath minute ventilation for control and *Hox5*$^{Dbx1Δ}$ mice in normal air at P50 (n = 5, ribbon = standard error).

H) Mean minute ventilation, shown as percent of sex-matched control littermates in normal air at P50 (*Hox5*$^{Dbx1Δ}$: 13.3 ± 13.5% increase, n = 5, p = 0.38).

I) Distribution plot of breath tidal volume for control and *Hox5*$^{Dbx1Δ}$ mice in 5% $CO_2$ at P50 (n = 5, ribbon = standard error).

J) Mean tidal volume, shown as percent of sex-matched control littermates in 5% $CO_2$ at P50 (*Hox5*$^{Dbx1Δ}$: 45.5 ± 4.7% decrease, n = 5, p = 0.00065).

K) Distribution plot of breath frequency for control and *Hox5*$^{Dbx1Δ}$ mice in 5% $CO_2$ at P50 (n = 5, ribbon = standard error).

L) Mean breath frequency, shown as percent of sex-matched control littermates in 5% $CO_2$ at P50 (*Hox5*$^{Dbx1Δ}$: 20.4 ± 12.3% increase, n = 5, p = 0.17).

M) Distribution plot of breath minute ventilation for control and *Hox5*$^{Dbx1Δ}$ mice in 5% $CO_2$ at P50 (n = 5, ribbon = standard error).

N) Mean minute ventilation, shown as percent of sex-matched control littermates in 5% $CO_2$ at P50 (*Hox5*$^{Dbx1Δ}$: 33.2 ± 9.8% decrease, n = 5, p = 0.027).

O) Percent change in tidal volume after exposure to 5% $CO_2$ at P50 (Control: 69.0 ± 5.0% increase, *Hox5*$^{Dbx1Δ}$: 25.6 ± 6.9% increase, n = 5, p = 0.0013).

P) Percent change in breath frequency after exposure to 5% $CO_2$ at P50 (Control: 44.7 ± 10.4% increase, *Hox5*$^{Dbx1Δ}$: 10.7 ± 6.0% increase, n = 5, p = 0.028).

Q) Percent change in minute ventilation after exposure to 5% $CO_2$ at P50 (Control: 149.1 ± 20.2% increase, *Hox5*$^{Dbx1\Delta}$: 39.8 ± 12.8% increase, n = 5, p = 0.0028).

**Figure S6. Phrenic MN activity is consistent within preparations**

A) Quiescence within bursts at P0, shown as percent of burst with no unit activity. Each point represents a single burst organized into columns for each sample (35-47 bursts, 5-7 samples).

B) Phrenic nerve activity during bursts at P0, shown as integrated burst activity normalized to burst duration and to the average of controls. Each point represents a single burst organized into columns for each sample (35-47 bursts, 5-7 samples).

C) Burst duration at P0, shown in seconds. Each point represents a single burst organized into columns for each sample (35-47 bursts, 5-7 samples).

**Figure S7. SST localization in the medulla**

A) Distance from the most caudal SST+ puncta to the spinal cord at P0 (Control: 616 ± 20 µm, *Hox5*$^{Dbx1\Delta}$: 240 ± 69 µm, n = 3-5, p = 0.025).

B) Distance from the most rostral SST+ puncta to moVII at P0 (Control: 416 ± 16 µm, *Hox5*$^{Dbx1\Delta}$: 413 ± 13 µm, n = 3-5, p = 1).

**Video S1. *Hox5*$^{Dbx1\Delta}$ pups experience respiratory distress**

Breathing behavior of a *Hox5*$^{Dbx1\Delta}$ pup and littermate control at P0.

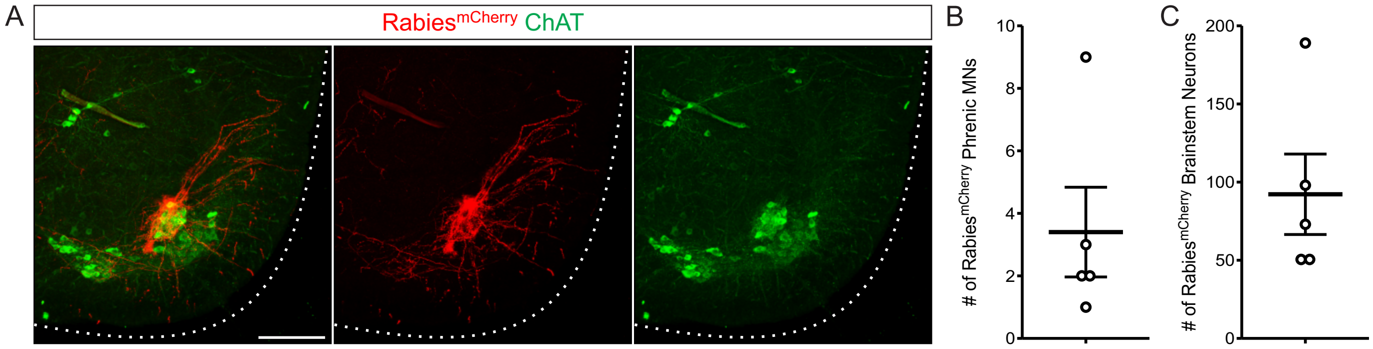
A
Rabies^mCherry ChAT
B
### of Rabies^mCherry Phrenic MNs
0
2
4
6
8
10
C
### of Rabies^mCherry Brainstem Neurons
0
50
100
150
200

Figure S1

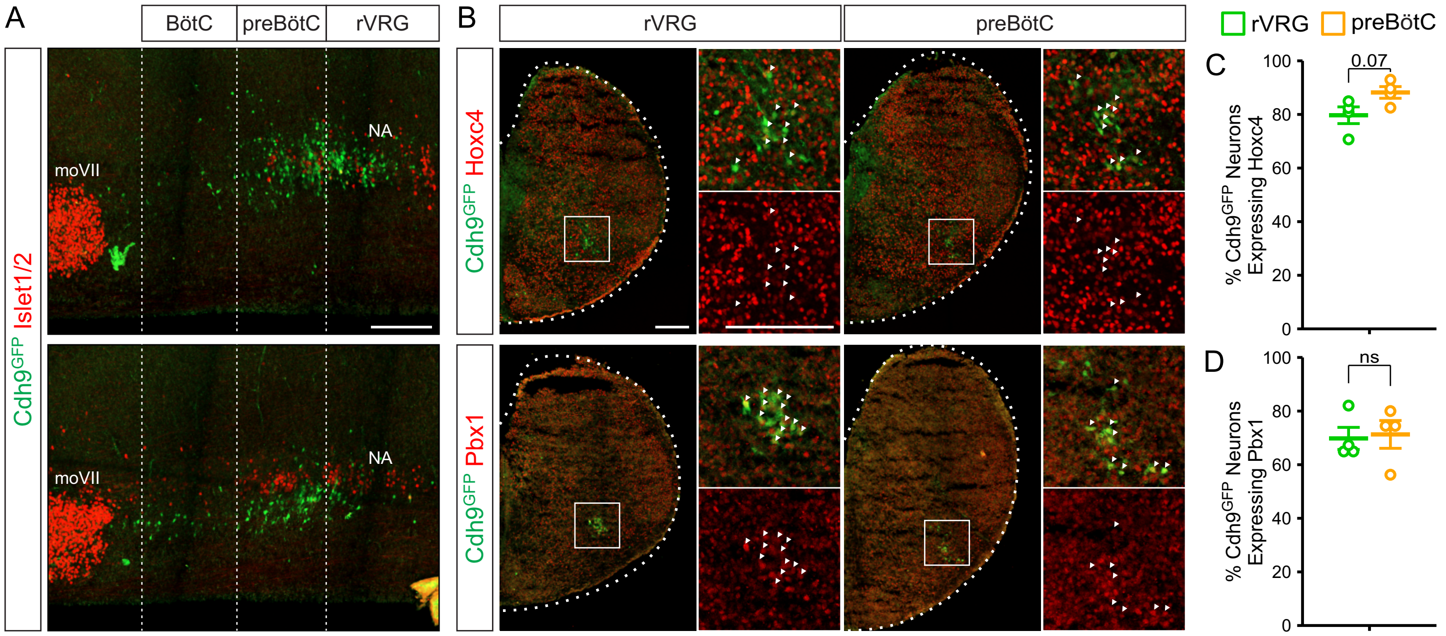
A
BötC
preBötC
rVRG
Cdh9GFP Islet1/2
moVII
NA
B
rVRG
preBötC
Cdh9GFP Hoxc4
Cdh9GFP Pbx1
rVRG
preBötC
C
0.07
% Cdh9GFP Neurons Expressing Hoxc4
D
ns
% Cdh9GFP Neurons Expressing Pbx1
0
20
40
60
80
100

Figure S2

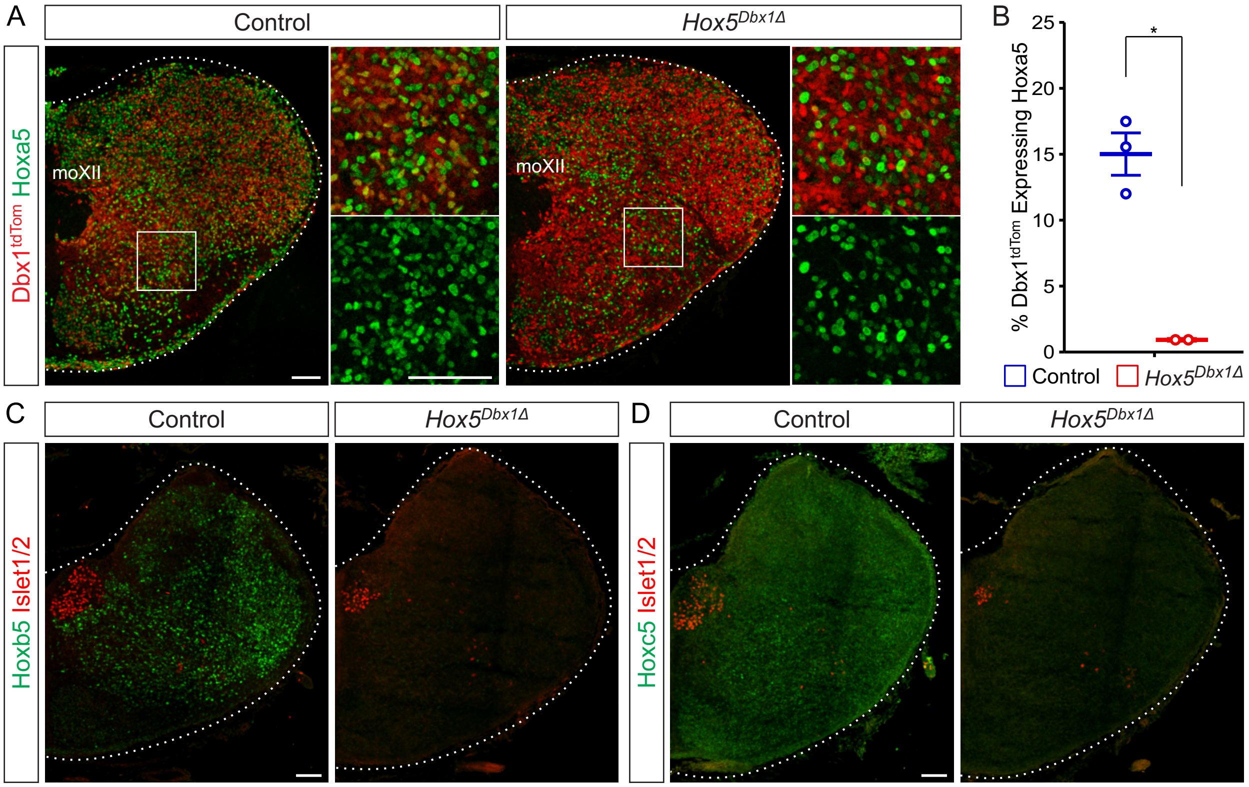
A
Control
$Hox5^{Dbx1\Delta}$
Dbx1tdTom Hoxa5
moXII
moXII
B
% Dbx1tdTom Expressing Hoxa5
25
20
15
10
5
0
*
Control
$Hox5^{Dbx1\Delta}$
C
Control
$Hox5^{Dbx1\Delta}$
Hoxb5 Islet1/2
D
Control
$Hox5^{Dbx1\Delta}$
Hoxc5 Islet1/2

Figure S3

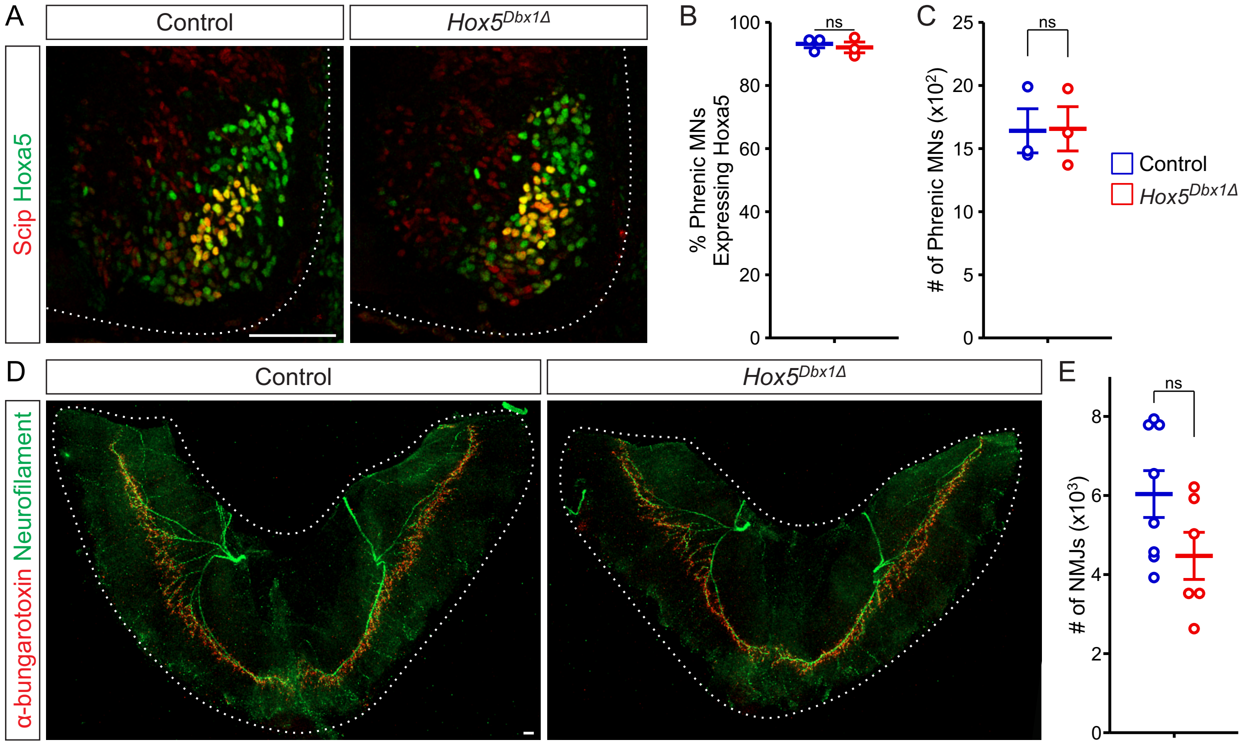
A
Control
Hox5Dbx1Δ
Scip Hoxa5
B
% Phrenic MNs Expressing Hoxa5
ns
C
### of Phrenic MNs (x10²)
ns
Control
Hox5Dbx1Δ
D
Control
Hox5Dbx1Δ
α-bungarotoxin Neurofilament
E
### of NMJs (x10³)
ns

Figure S4

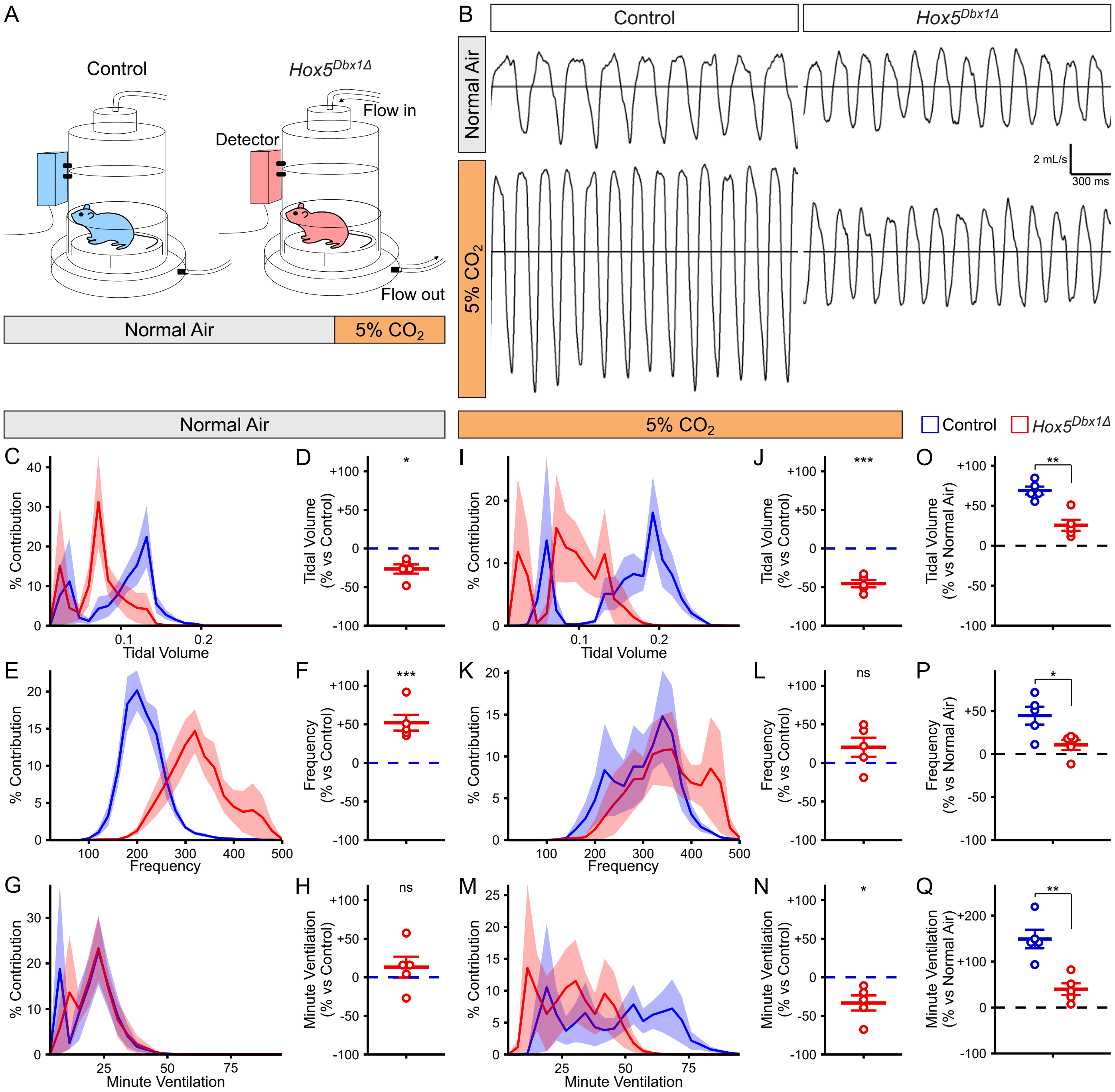
A
Control
Hox5Dbx1Δ
Flow in
Detector
Flow out
Normal Air
5% CO2
B
Control
Hox5Dbx1Δ
Normal Air
5% CO2
2 mL/s
300 ms
Normal Air
5% CO2
Control
Hox5Dbx1Δ
C
% Contribution
Tidal Volume
D
Tidal Volume (% vs Control)
*
E
% Contribution
Frequency
F
Frequency (% vs Control)
***
G
% Contribution
Minute Ventilation
H
Minute Ventilation (% vs Control)
ns
I
% Contribution
Tidal Volume
J
Tidal Volume (% vs Control)
***
K
% Contribution
Frequency
L
Frequency (% vs Control)
ns
M
% Contribution
Minute Ventilation
N
Minute Ventilation (% vs Control)
*
O
Tidal Volume (% vs Normal Air)
**
P
Frequency (% vs Normal Air)
*
Q
Minute Ventilation (% vs Normal Air)
**

Figure S5

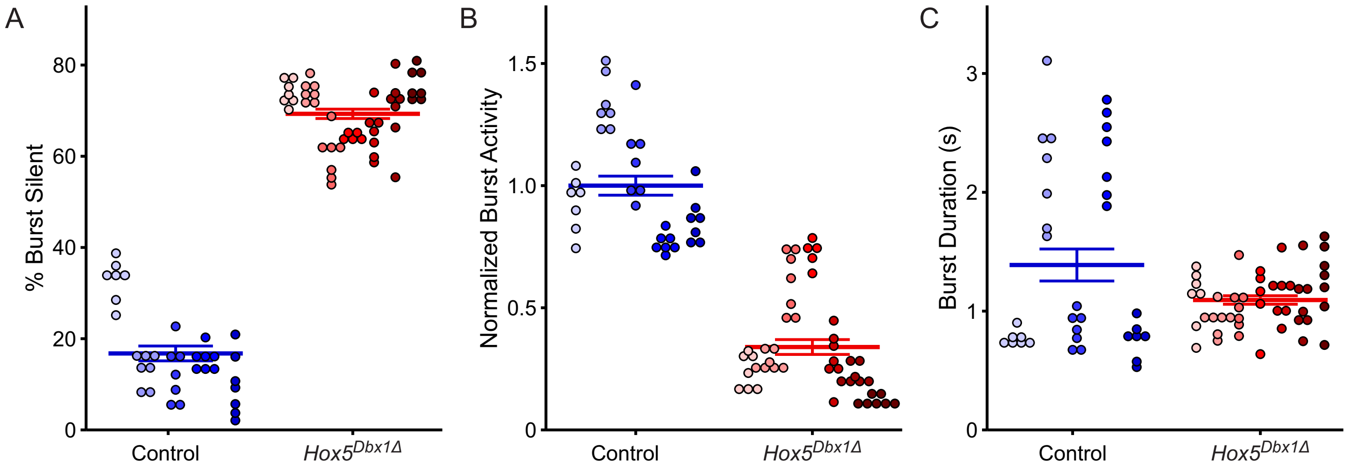
A
% Burst Silent
0
20
40
60
80
Control
Hox5Dbx1Δ
B
Normalized Burst Activity
0
0.5
1.0
1.5
Control
Hox5Dbx1Δ
C
Burst Duration (s)
0
1
2
3
Control
Hox5Dbx1Δ

Figure S6

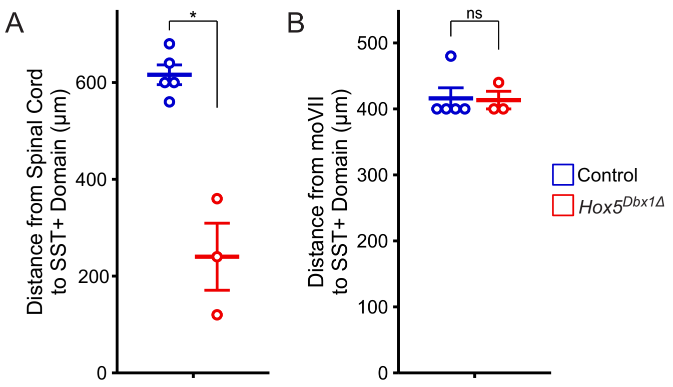
A
B
*
ns
Distance from Spinal Cord
to SST+ Domain (µm)
Distance from moVII
to SST+ Domain (µm)
0
200
400
600
0
100
200
300
400
500
Control
Hox5Dbx1Δ

Figure S7
